# Quantitative genetics of photosynthetic trait variation in maize

**DOI:** 10.1101/2024.11.25.625283

**Authors:** Waqar Ali, Marcin Grzybowski, J. Vladimir Torres-Rodríguez, Fangyi Li, Nikee Shrestha, Ramesh Kanna Mathivanan, Gabriel de Bernardeaux, Khang Hoang, Ravi V. Mural, Rebecca L. Roston, James C. Schnable, Seema Sahay

## Abstract

Natural genetic variation in photosynthesis-related traits can aid both in identifying genes involved in regulating photosynthetic processes and developing crops with improved productivity and photosynthetic efficiency. However, rapidly fluctuating environmental parameters create challenges for measuring photosynthetic parameters in large populations under field conditions. We measured chlorophyll fluorescence and absorbance-based photosynthetic traits in a maize diversity panel in the field using an experimental design that allowed us to estimate and control multiple confounding factors. Controlling the impact of day of measurement and light intensity as well as patterns of two-dimensional spatial variation in the field substantially increased heritability with the heritability of 7 out of 14 traits measured exceeding 0.4. We were able to identify high confidence GWAS signals associated with variation in four spatially corrected traits (the quantum yield of photosystem II, non-photochemical quenching, redox state of Q_A_, and relative chlorophyll content). Insertion alleles for *Arabidopsis* orthologs of three candidate genes exhibited phenotypes consistent with our GWAS results. Collectively these results illustrate the potential of applying best practices from quantitative genetics research to address outstanding questions in plant physiology and understand the mechanisms underlying natural variation in photosynthesis.

**Highlights:**

● Controlling spatial and environmental confounding factors increased heritability of photosynthetic traits.
● GWAS identified high confidence signals associated with variation in relative chlorophyll, ΦPSII, ΦNPQ, and qL.
● Insertion alleles of the *Arabidopsis* orthologs of maize candidate genes exhibited photosynthesis related phenotypes consistent with the GWAS results.

## Introduction

Ongoing and forecasted climate changes are placing a growing strain on global food security and make ongoing increases in crop resilience and productivity essential. Improvements in photosynthetic efficiency have the potential to significantly increase yields (Long *et al*., 2015). Photosynthesis, the biological process of converting light energy and atmospheric CO_2_ into chemical energy and organic compounds, is a vital process for the sustainability of food production systems in the world. A better understanding of the photosynthetic processes may help to identify target traits and target genes for improving inherent photosynthetic efficiency to further enhance productivity without expanding land use, increased water consumption, or requiring further growth in nitrogen fertilizer applications. A number of proof-of-concept studies have demonstrated improvements in photosynthetic efficiency and productivity in crop plants through transgenic approaches (Kromdijk *et al*., 2016; Driever *et al*., 2017; López-Calcagno *et al*., 2019; De Souza *et al*., 2022). Genetic engineering-based approaches that have demonstrated potential to increase photosynthetic productivity include the manipulation of various photosynthetic traits such as non-photochemical quenching, mesophyll conductance, rubisco activity, and sedoheptulose-bisphosphatase via overexpression to increase photoprotection, water-use-efficiency, CO2 fixation, and stomata regulation which led to improve photosynthetic efficiency and yield in tobacco, *Arabidopsis*, and rice (Kromdijk *et al*., 2016; Simkin *et al*., 2017; Hubbart *et al*., 2018; South *et al*., 2019).

It may also be possible to exploit natural genetic variation for photosynthetic parameters in crop plants either to identify targets of genetic engineering to improve photosynthetic efficiency (Gu *et al*., 2014; Basu *et al*., 2019; Faralli and Lawson, 2020; Acevedo-Siaca *et al*., 2021; Meena *et al*., 2021; Sahay *et al*., 2023, 2024) or to optimize photosynthesis capacity via marker-assisted selection or genomic selection (van Bezouw *et al*., 2019; Theeuwen *et al*., 2022). It has been suggested that exploiting variations in multiple photosynthetic traits and aggregating them could produce synergistic impacts on crop yields (Sinclair *et al*., 2019; Araus *et al*., 2021). However, as photosynthetic performance varies in response to light intensity, temperature, access to water, and other stresses as well as in response to diurnal and circadian cycles, quantifying the impacts of natural variants for the large populations used for quantitative genetic investigation or crop improvement can be challenging under field conditions.

Rapid fluorescence-based approaches to measure photosynthetic traits have been employed in high throughput forward genetic screens under controlled conditions to identify a number of the genes encoding core components of the photosynthetic regulatory apparatus in algae and the model plant *Arabidopsis thaliana* (Niyogi *et al*., 1997, 1998; Baker, 2008). More recently, several field portable technologies have become available for rapidly estimating photosynthesis related parameters using a combination of fluorescence and absorbance based technologies. Previous efforts to use these technologies to identify genes controlling natural variation in photosynthetic parameters have been limited by the the low heritability of photosynthetic traits measured under field conditions (Dramadri *et al*., 2021; Liu *et al*., 2023). However, these previous attempts did not directly control for either the impact of changes in environmental factors during data collection on the traits of interest nor model and control for 2D spatial variation within individual field experiments.

Here, we evaluated the potential of incorporating statistical controls for the impact of fluctuating environmental conditions to enable the identification of genes controlling variation in photosynthesis related traits under field conditions using low-cost, field portable tools. We collected data from a large replicated maize diversity panel in the field employing an experimental design that allowed us to separately estimate the impact of multiple confounding environmental factors. Consistent with previous reports, the photosynthesis related traits scored largely exhibited extremely low heritabilities (e.g. the proportion of total variance attributable to differences between genotypes) in our initial model, but controlling for confounding environmental factors allowed many more traits to reach levels of heritability suitable for directed breeding efforts via genomic prediction or gene discovery via genome-wide association studies (GWAS). We identified well-supported trait-associated markers in GWAS for four of the fourteen traits initially profiled. In three cases, insertion alleles of the *Arabidopsis* orthologs of maize candidate genes also exhibited photosynthesis related phenotypes consistent with the maize GWAS results. However, further in-depth characterization efforts will be necessary to confirm the roles of the candidate genes identified in determining variation in photosynthetic traits under field conditions.

## Materials and Methods

### Maize association panel field experiment

A subset of 752 maize (*Zea mays* L.) genotypes from the Wisconsin Diversity Panel (Mazaheri *et al*., 2019) were grown in two randomized blocks of 840 plots for a total 1,680 plots consisting of one entry of each unique genotype per block with remaining plots filled with a single repeated check at the University of Nebraska–Lincoln’s Havelock Farm (40.852N, 96.616W). A list of the specific genotypes employed in our study is provided in supplementary data files. Each plot consisted of two parallel rows of plants from a single genotype with 30 inches (0.76 m) separating the two rows. The rows ran from east to west and individual plots were positioned in a 60 East-West х 28 North-South grid. The experimental field was surrounded by an additional border of plots planted with a single uniform genotype to mitigate edge effects. Final plant density varied due to differences in germination and plant survival rates, as well as the removal of any off-type plants observed prior to sampling. The seeds were sown using a four row Almaco plot planter on the 6^th^ of May 2020. Nitrogen fertilizer was applied via a single urea ammonium nitrate prior to planting. Weeds were controlled by a pre-emergence application of atrazine (Syngenta) according to the manufacturer’s recommendations, followed by manual weed removal throughout the growing season. No irrigation was provided prior to or during the growing season. The layout and the details of the field setup have been previously described (Mural *et al*., 2022; Sun *et al*., 2022; Sahay *et al*., 2023).

### Scoring fluorescence– and absorbance-based photosynthesis-related traits

Fluorescence– and absorbance-based photosynthetic-related traits were scored using a set of six field portable spectrophotometers (MultiSpeQ V2.0; PhotosynQ, East Lansing, MI, USA; (Kuhlgert *et al*., 2016) between 9:30 am to 2:30 pm on July 23^rd^ (1,397 plots phenotyped), 24^th^ (1,665 plots phenotyped), 25^th^ (280 plots phenotyped), and 28^th^ (1,679 plots phenotyped). All the measurements were performed using the Pulse-Amplitude-Modulation (PAM) method as implemented using Photosynthesis Rides protocol provided by the manufacturer. The 14 traits quantified and analyzed as part of this study were chlorophyll SPAD (relative chlorophyll content), light adapted maximum efficiency of PSII in the light (Fv’/Fm’), redox state of quinone A; Q_A_ (qL), total electrochromatic shift (ECS_T_), proton conductivity (gH^+^), steady-state proton flux (vH^+^), transient values of theoretical non-photochemical quenching (NPQ_T_), quantum yields of photosystem II (ΦPSII), non-photochemical quenching (ΦNPQ), and other non-regulated energy dissipation (ΦNO), photosystem I (PSI) active centers (PSI-ac), PSI-open centers (PSI-opc), PSI-oxidized centers (PSI-oxc), and PSI-over reduced centers (PSI-orc). Each plot, excluding those without any healthy plants present, was measured three times, with each measurement occurring on a different day to avoid confounding any potential date-of-sampling effect with genotype effects. For each measurement, a plant within the plot was arbitrarily selected by the sampler, avoiding edge plants when possible. All measurements were performed on a fully expanded leaf, targeting a point midway between the ligule and the leaf tip. The field portable spectrophotometers automatically recorded the specific time, date, intensity of photosynthetically active radiation, ambient temperature and ambient humidity data during each measurement.

### Phenotypic data processing and quality control

Initial heritability estimates were generated for each trait using plot level average for each trait. These plot level averages were calculated after dropping individual measurements with extreme values outside of the expected range for each trait see cutoffs in Table S1. Heritability was defined via the equation H^2^ = Vg/(V_g_+V_ε_/N) where V_g_ and V_ε_ are the variance attributable to genotype and residual variance respectively. Both parameters were estimated by fitting the model Trait ∼ Genotype + Residual implemented using the lme4 package in R (Bates *et al*., 2015). Estimates of the proportion of phenotypic variance explained by genetic and non-genetic factors were generated using lme4, fitting the model Trait ∼ Genotype + Row + Column + Day + Light Intensity, with all variables treated as random effects and dividing the variance attributed to each factor for a given phenotype by the total variance for that phenotype.

Final heritability estimates were generated using the same approach as the initial heritability measurements using corrected plot-level values generated in SpATS rather than the simple plot level averages employed for the initial estimates. Corrected plot level values were generated in SpATS v1.0-18 (Rodríguez-Álvarez *et al*., 2018) fitting light intensity and ambient temperature as a fixed effect, plotID and day as random effects and, modeling 2D spatial effects on each phenotype using 15 column knots and 31 row knots. Best linear unbiased estimates were generated as described above for corrected plot level values with the modification that genotype was substituted for plotID in SpATS (Fig. S1).

### Genome-wide association studies

Genome-wide association studies (GWAS) were conducted using the FarmCPU algorithm as implemented in the rMVP package (version, 1.0.8) (Liu *et al*., 2016; Yin *et al*., 2021) including three PCs calculated from genetic marker data as additional covariates. The resequencing-based marker set generated in (Grzybowski *et al*., 2023) was subset to only those biallelic markers which exceeded a minor allele frequency of 0.05 considering only those individual represented in this study and excluding markers where >20% of genotypes were classified as heterozygous, resulting in a set of 11.8 million genetic markers. The effective number of independent markers represented by this dataset was estimated to be 4.7 million via GEC 0.2 (Li *et al*., 2012). For each trait, the FarmCPU GWAS analysis was conducted 100 times, with 10% of trait values masked for each iteration and a p.threshold setting of 1.06 x 10^-8^, calculated based on the Bonferroni correction applied to an initial *p*-value of 0.05 and the estimated number of independent markers. Markers were considered to be significant in a given interaction if they exceeded a *p*-value threshold of 1.06 x 10^-8^. Resampling model inclusion probabilities (RMIPs) were calculated by dividing the number of iterations in which that marker was significantly associated with a given trait (*p* < 1.06 x 10^-8^) by the total number of iterations run (100). Linkage disequilibrium between markers in mapping intervals was estimated using PLINK 1.9 (Purcell *et al*., 2007).

### Chlorophyll ground truth data

Absolute chlorophyll content was scored separately from the measurements described above, with data collected from a single leaf of a single plant from each of 318 plots using a handheld chlorophyll meter (MC-100, Apogee Instruments, Inc., Logan, UT). Absolute chlorophyll measurements were collected on nine days spanning a thirteen day sampling period between July 8 to July 20, 2020 (see (Tross *et al*., 2024).

### eQTL analysis

The eQTL results presented in this study were conducted using gene expression data of 693 genotypes profiled using mRNA-seq of RNA samples extracted from mature leaf tissue collected in a two hour period on July 8^th^ 2020 (Torres-Rodríguez *et al*., 2024). QTL mapping was performed using the mixed linear method as implemented within rMVP package (1.0.8) in R (Price *et al*., 2006; Yin *et al*., 2021) and the same set of 11.8M segregating markers employed for GWAS above. Three PCs and kinship matrix calculated from genetic marker data were included as covariates when conducting genome wide association studies for gene expression levels. Associations between markers and gene expression were considered significant if they exceeded a *p*-value threshold of 1.06 x10^-8^.

### Characterization of Arabidopsis mutants

T-DNA insertion lines were tested for 3 genes: SALK_007055, carrying an insertion in AT1G10500 (*ATCPISCA* or *Chloroplast-localized ISCA-like protein*), SALK_080503, carrying an insertion in AT2G26660 (*ATSPX2* or *SPX domain gene 2*), and SALK_047115, carrying an insertion in AT5G42520 (*ATBPC6* for *Basic Pentacysteine 6*). Homozygosity of each T-DNA insertion within individual plants from each stock was confirmed by PCR, using the T-DNA-specific primer LBb1.3 along with gene-specific left and right primer designed through the Salk T-DNA primer design tool (http://signal.Salk.edu/tdnaprimers.2.html) and detailed in (Table S2).

Seeds for all T-DNA insertional mutants and their wild-type control (Columbia ecotype, Col-0, CS6000) were obtained from the Arabidopsis Biological Resource Center (ABRC) at Ohio State University (Alonso *et al*., 2003). Seeds were stratified for 4 days at 4°C in the dark, then sown in 3.5 inch × 3.5 inch pots (SQN03500B66, Hummert International, Earth City, MO, USA) filled with soil-less potting mix (1220338; BM2 Germination and Propagation Mix; Berger, SaintModeste, Canada). Pots were placed in trays (6569630; Hummert International) filled with 2 centimeters of water at the bottom and covered with a clear plastic dome (65696400; Hummert International) until germination. The trays were placed in a reach-in growth chamber (AR-66 L2; Percival, Perry, IA, USA) at 21°C day/18°C night with a 10-hour-light/14-hour-dark photoperiod (200 μmol m^−2^ sec^−1^) and 60% relative humidity. One week after germination, seedlings were thinned to one per pot; plants were watered and repositioned at random locations in the chamber three times a week.

Relative chlorophyll, ΦPSII, and ΦNPQ were measured using MultispeQs with the same protocol employed for field measurement. The initial set of low light measurements were taken on four-week-old plants under low-light conditions (LL; 200 μmol m^−2^ sec^−1^). High light measurements were collected from the same plants after the plants were moved to a high-light treatment (HL; 550 μmol m^−2^ sec^−1^ at 24 °C/22 °C day/night with an 8 hour day/16 hour night) in a walk-in growth chamber (Conviron model GR48, Controlled Environments, Manitoba, Canada) for 24 hours.

## Statistical analysis

Statistical analyses of photosynthetic parameter measurements in *Arabidopsis* mutants were performed using R software v.4.3.2, with the package lme4. Difference between mutant and wild type were tested with unpaired, two-tailed t-test.

## Results

### Repeatability of field-scored photosynthetic traits is enhanced by removing outliers and controlling for spatial variation

Considering simple plot level averages for the three independent measurements collected per plot, with one exception, all photosynthesis-related traits assayed in this experiment exhibited heritability of < 0.25 (Fig. 1A). The sole exception was relative chlorophyll content with an estimated heritability of 0.61. Excluding photosystem I (PSI) active centers (PSI-ac), all PSI-related traits exhibited heritabilities less than 0.1. The heritabilities for photosynthesis-related traits were substantially lower than those observed for whole plant phenotypes measured in the same field experiment. The mean and median heritability of 28 manually measured plant phenotypes collected from the same field (Mural *et al*., 2022) were 0.75 and 0.85 respectively.

**Fig. 1.**
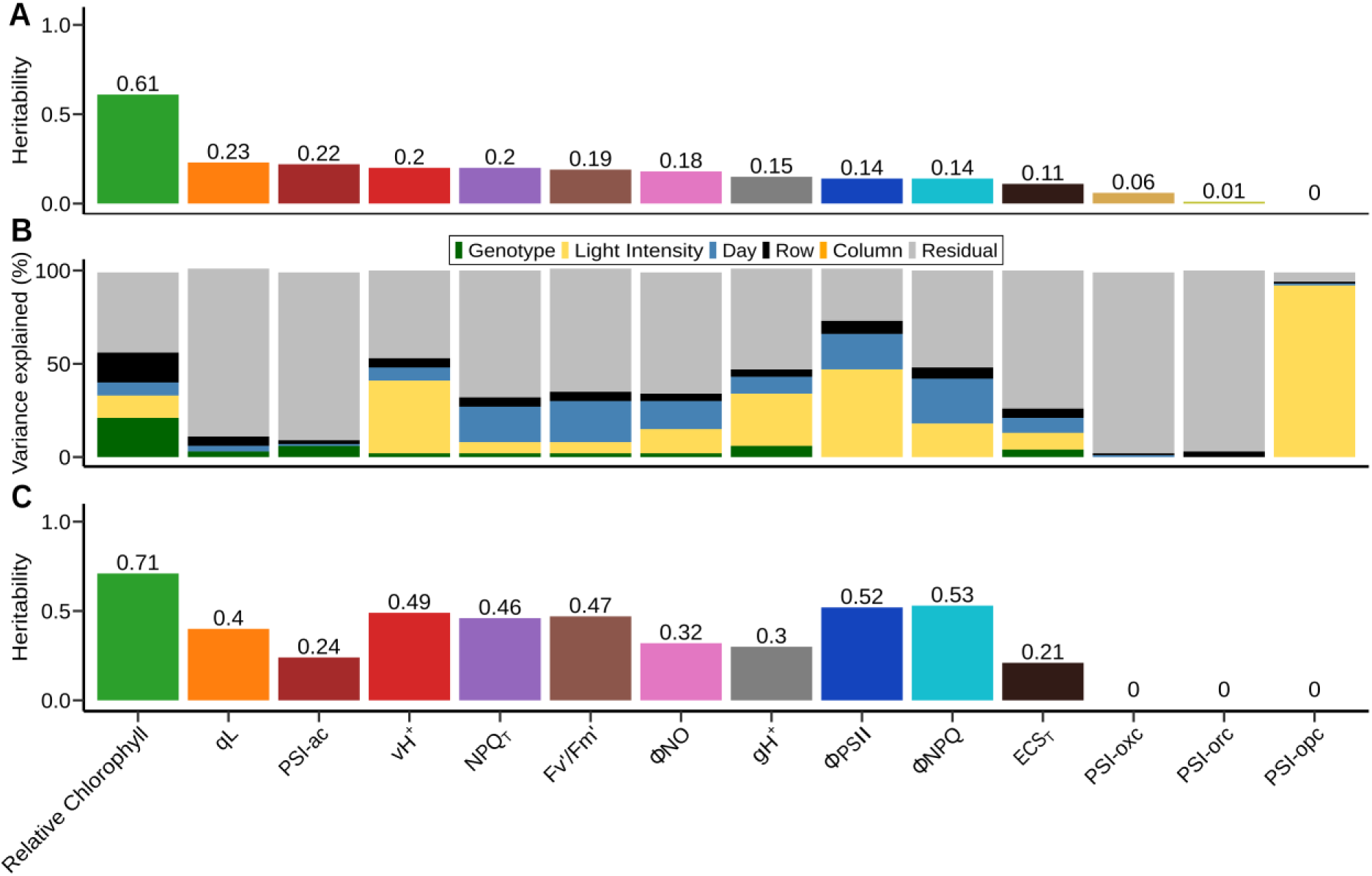
Factors explaining variation in measured values for photosynthesis-related traits under field conditions. **(A)** Estimated genotype-level heritability across two replicated plots in the same field for plot-level averages, uncorrected for environmental or spatial confounders, of fourteen traits scored in the field using MultiSpeQs. **(B)** Estimated proportion of variance in individual measurements of the same fourteen traits attributable to genotype, light intensity recorded at the time of collection, day in which the measurement was taken, and row and column position within the field. **(C)** Estimated genotype-level heritability across two replicated plots in the same field for plot level trait estimates generated after correcting for the effects of light intensity and day as well as 2D spatial variation throughout the field using SpATS.

The distribution of values recorded for many of the photosynthesis-related traits were significantly different across the four days over which data collection occurred. Day of collection explained 7% variation in relative chlorophyll, 19% in quantum yield of photosystem II (ΦPSII) and 24% in non-photochemical quenching (ΦNPQ) of total variance (Fig. 1B, Table S3). Light intensity varied substantially over the time period where data was collected, with several prolonged periods of lower intensity light resulting from clouds obscuring the sun (Fig. S2). Variation in light intensity at the time of collection also explained a percentage of variance ranging from 12 to 47% of the total variation for relative chlorophyll, ECS_T_, gH^+^, vH^+^, ΦPSII, and explaining 92% of the variation in PSI open centers (PSI-opc), a trait with essentially no heritability between replicates of the same genotype in different parts of the field (Fig. 1B, Table S3). Sixteen (16%) of the variance in measured relative chlorophyll content was explainable by differences between different rows in the field. It should be noted that this is likely an underestimate of the total variation in these traits explained by with-in field spatial variation as many factors with the potential to influence plant phenotypes –-e.g. nutrient availability, soil type, soil moisture, elevation –-will vary along gradients not easily captured by either row or column variables.

Controlling for the impact of light intensity and day as well as patterns of two-dimensional spatial variation in traits across the research field increased the heritability of corrected plot-level estimates for thirteen out of fourteen phenotypes (Fig. 1C). ΦPSII and ΦNPQ, estimates of the proportions of light captured by photosystem II which are employed for productive photochemistry and dissipated via photoprotection mechanisms respectively, were the largest increases in heritability. The heritability of ΦPSII increased from 0.14 when using simple average values to 0.52 when correcting for spatial variation and confounding environmental factors. Similarly, the heritability of ΦNPQ increased from 0.14 to 0.53. Traits related to photosystem I exhibited the smallest increases in heritability and included the only traits where correction for confounding factors either failed to increase heritability (PS1-opc) or decreased heritability (PS1-orc).

### Genetic loci linked to variations in photosynthesis-related traits

Genetic markers could be linked to variation in four of the fourteen traits measured in this study at an RMIP (the proportion of times a marker is included in the resampling-data based on iterations in the FarmCPU model) threshold of > 0.2, including relative chlorophyll content (one hit), ΦPSII (two hits), ΦNPQ (two hits), and qL (one hit) (Fig. 2A, Table 1).

**Fig. 2.**
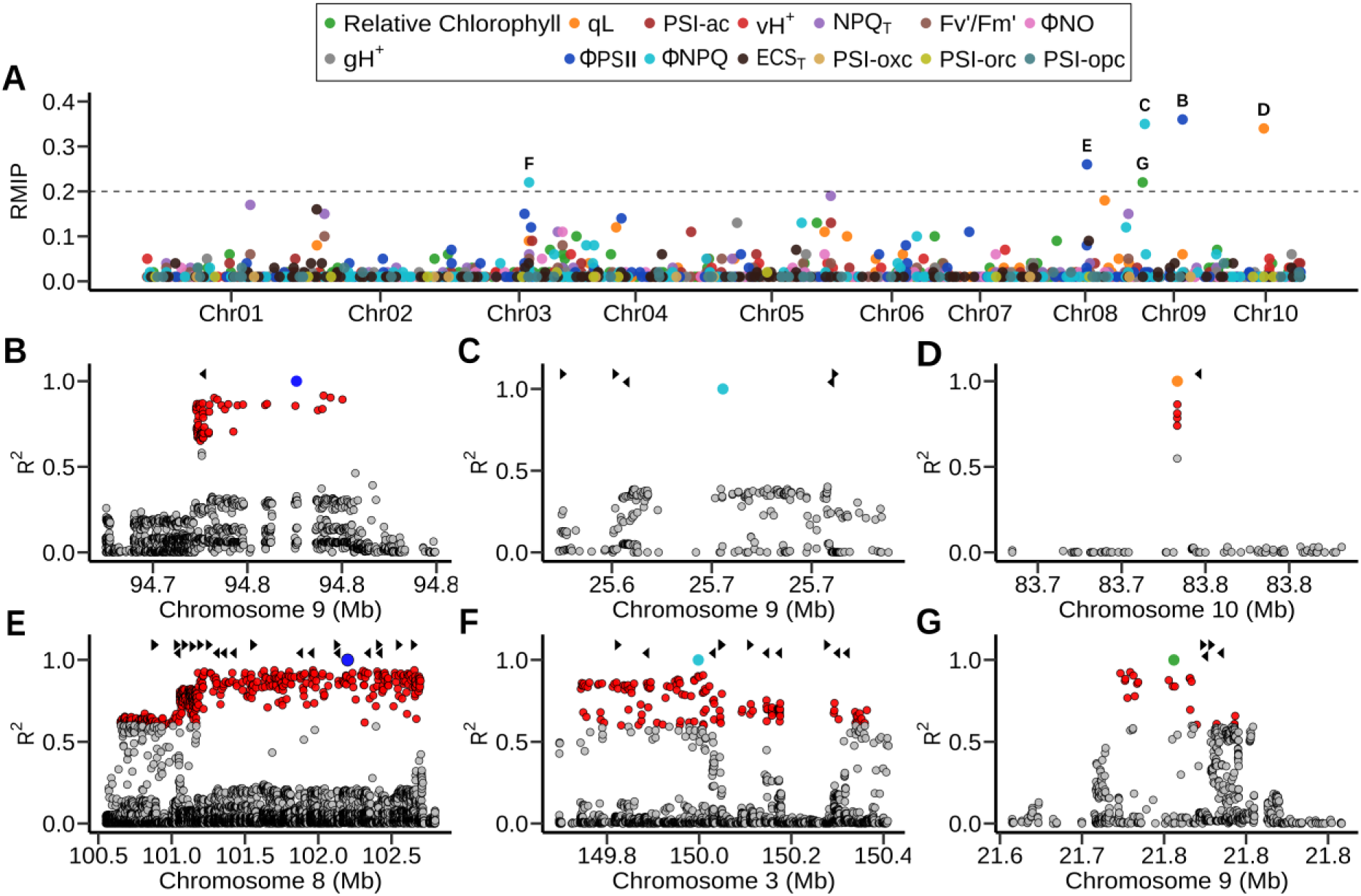
Genetic markers in the maize genome significantly associated with variation in photosynthetic traits under field conditions. **(A)** Statistical support and position on the maize genome for individual genetic markers associated with variation in fourteen photosynthetic traits scored for maize genotypes from the Wisconsin Diversity panel under field conditions in 2020. Position on the x-axis indicates the physical position of the marker on the B73 RefGen V5 genome assembly. Position on the y-axis indicates the proportion of 100 iterations of FarmCPU GWAS where the marker was significantly associated with the given trait. Dashed line indicates a RMIP threshold of > 0.2. Letters indicate the figure panel where further detail is provided for individual markers identified above this threshold. **(B)** Region containing markers in elevated linkage disequilibrium with a genetic marker associated with variation in ϕPSII located at position 94,775,951 on chromosome 9. Circles indicate the positions (x-axis) and linkage disequilibrium with the trait associated marker (y-axis) for other genetic markers in this interval. To capture the context of linkage disequilibrium surrounding the interval of interest, the range shown begins approximately 4 kilobases before the first marker in linkage disequilibrium >0.6 with the trait associated marker (red circles) and extends approximately four kilobases beyond the last marker in linkage disequilibrium with the trait associated marker. Black triangles indicate the positions and strands of annotated gene models within this interval. **(C)** Region containing markers in elevated linkage disequilibrium with a genetic marker associated with variation in ϕNPQ located at position 25,683,346 on chromosome 9. All figure elements are as defined in panel B. **(D)** Region containing markers in elevated linkage disequilibrium with a genetic marker associated with variation in qL located at position 83,741,616 on chromosome 10. All figure elements are as defined in panel B. **(E)** Region containing markers in elevated linkage disequilibrium with a genetic marker associated with variation in ϕPSII located at position 102,200,025 on chromosome 8. All figure elements are as defined in panel B. **(F)** Region containing markers in elevated linkage disequilibrium with a genetic marker associated with variation in ϕNPQ located at position 83,741,616 on chromosome 3. All figure elements are as defined in panel B. **(G)** Region containing markers in elevated linkage disequilibrium with a genetic marker associated with variation in relative chlorophyll located at position 21,755,970 on chromosome 9. All figure elements are as defined in panel B.

**Table 1.** Genomic regions and trait association summary from genome wide association studies.

| **CHR** | **POS** | **Trait** | **RMIP** | **Associated Genomic Interval** | **Number of Genes** | **Other Associations** |
| --- | --- | --- | --- | --- | --- | --- |
| Chr9 | 94775951 | ΦPSII | 0.36 | 94,725,951 - 94,825,951, | 1 | ΦNPQ, 0.02 |
| Chr9 | 25683346 | ΦNPQ | 0.35 | 25,633,346 - 25,733,346 | 5 | - |
| Chr10 | 83741616 | qL | 0.34 | 83,691,616 - 83,791,616 | 1 | Fv'/Fm', 0.01 |
| Chr8 | 102200025 | ΦPSII | 0.26 | 102,150,025, - 102,250,025 | 22 | - |
| Chr3 | 149998190 | ΦNPQ | 0.22 | 149,948,190 - 150,048,190 | 12 | - |
| Chr9 | 21755970 | Relative Chlorophyll | 0.22 | 21,705,970-21,805,970 | 4 | - |

These four traits included the three traits with the highest heritability among those examined in this study after correcting for spatial variation and other confounding factors. A parallel study conducted including another statistical model which also sought to account for device-to-device variation in measurement, a rough proxy for identity of the person making measurements and selecting which plants to measure, provided roughly equivalent results (Fig. S3).

Between 1 and 22 annotated gene models were close enough to the individual trait associated markers to be considered plausible candidate genes using a linkage disequilibrium (R^2^) threshold of 0.6 (Table 1). The signal on chromosome 9 associated with variation in ΦPSII was located within the gene body of (Zm00001eb386270) (Fig. 2B, Table S4) an SPX domain-containing membrane protein which has been referred to as either *SPX6* (Xiao *et al*., 2021) or *SPX1* (Luo *et al*., 2024). Below, to avoid confusion with the Arabidopsis orthologs, also assigned numerical SPX designations, we will refer to this gene as *ZmSPX2*. The signal on chromosome 9 associated with variation in ΦNPQ was in an interval containing five annotated genes including a peptidyl-prolyl cis-trans isomerase (Zm00001eb378260) and *bbr4* (Zm00001eb378270), a member of the BPC plant-specific transcription factor family (Fig. 2C, Table S4). The signal on chromosome 10 associated with variation in qL specifically tagged an MYB transcription factor *(ZmMYB26*; Zm00001eb416530) annotated as playing a role in leaf senescence (Fig. 2D, Table S4). The signal on chromosome 8 associated with variation in ΦPSII was located in a region of elevated linkage disequilibrium including 22 annotated genes (Fig. 2E, Table S4). The signal on chromosome 3 associated with variation in ΦNPQ was also located in a region of elevated linkage disequilibrium with 12 annotated gene models located within the plausible interval around the trait associated SNP (Fig. 2F, Table S4). The final marker-trait association which exceeded a RMIP threshold of 0.2 was a marker located at approximately 22 megabases on chromosome 9 associated with variation in relative chlorophyll content. The LD window surrounding this gene included a cluster of four annotated gene models (Fig. 2G, Table S4).

The gene model, Zm00001eb377130, closest to the trait associated marker for relative chlorophyll on chromosome 9 encodes a chloroplast localized protein involved in iron-sulfur metabolism (Fig. 3A). The SNP associated with variation in relative chlorophyll content assessed using the method employed in this study (Fig. 3B) was also significantly predictive of absolute chlorophyll measured at a different time point for a subset of plots using an independent methodology (see methods) (Fig. 3C). The expression of Zm00001eb377130 in mature leaf tissue is associated with a large effect cis-eQTL with the genetic markers most significantly associated with the gene located immediately downstream of the annotated gene model (Fig. 3A). The genetic marker associated with variation in relative chlorophyll content is also significantly associated with variation in the expression of Zm00001eb377130 (Fig. 3D).

**Fig. 3.**
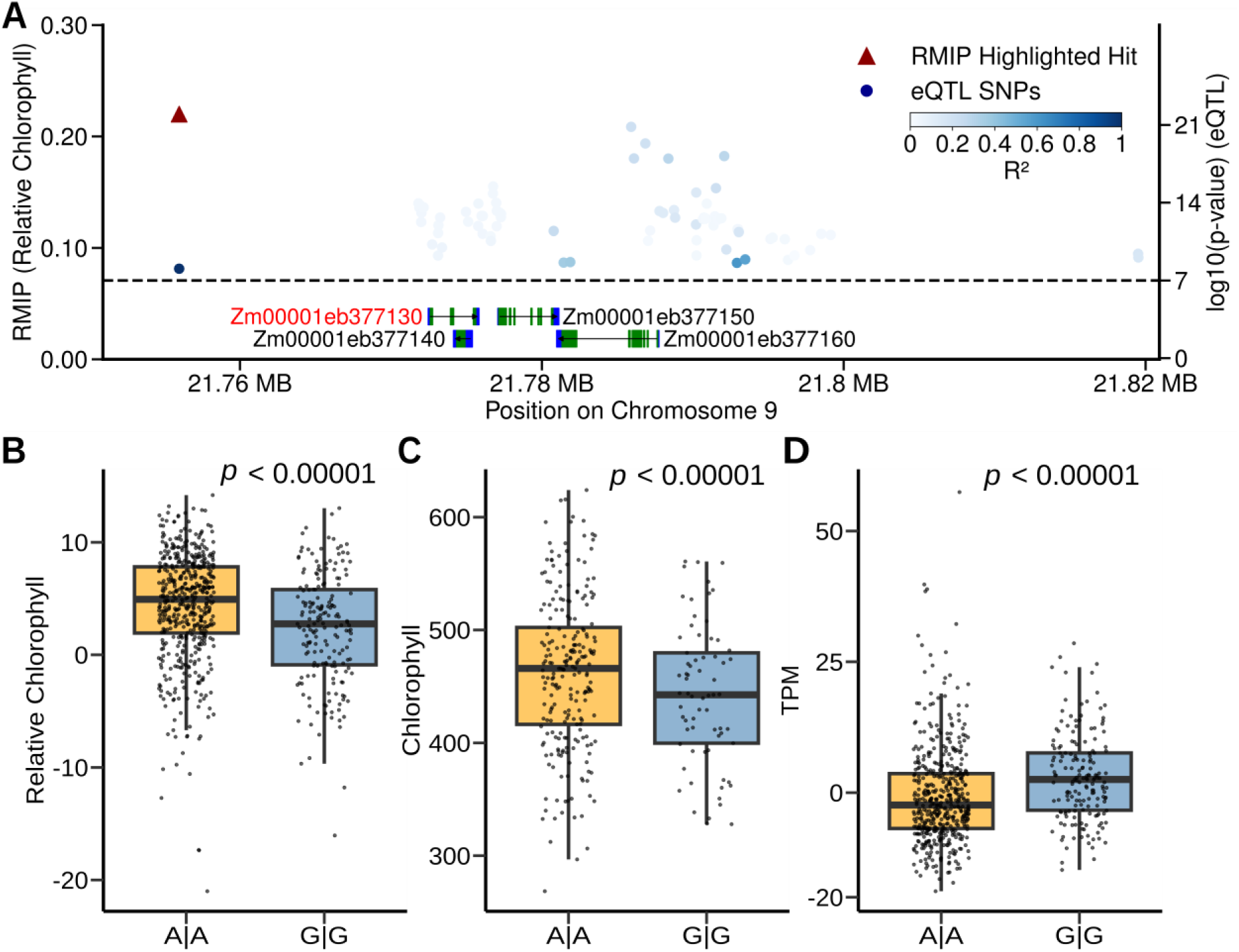
A genetic marker associated with variation in relative chlorophyll content is also associated with variation in absolute chlorophyll content and the expression of a nearby gene encoding a plastid localized protein. **(A)** Region containing both the genetic marker associated with relative chlorophyll content (Chr09:21,755,970, red triangle) and the four genes contained within the linkage disequilibrium defined interval surrounding this marker. Black arrows indicate the strand of genes, green boxes the positions of protein coding exons and blue boxes the positions of 5’ and 3’ UTRs. Blue circles indicate the positions (x-axis) of genetic markers significantly associated with variation in the expression of Zm00001eb377130 and statistical significance of that association (y-axis). Color of circles indicates the degree of linkage disequilibrium between markers associated with variation in gene expression and Chr09:21,755,970. **(B)** Difference in relative chlorophyll content between maize genotypes homozygous for the major (A|A) (n=519) and minor (G|G) alleles (n=183) of Chr09:21,755,970. The *p*-value shown was calculated via two sample t-test implemented in R. **(C)** Difference in absolute chlorophyll content scored for a subset of maize genotypes scored using a handheld chlorophyll meter taken from (Tross *et al*., 2024). Genotypes homozygous for the major (A|A) alleles (n=178). Genotypes homozygous for the minor (G|G) allele (n=66). **(D)** Difference expression of Zm00001eb377130 in mature leaf tissue between maize genotypes homozygous for the major (A|A) alleles (n=480) and minor (G|G) alleles (n=171) of Chr09:21,755,970. The *p*-value shown was calculated via a two sample t-test as implemented in R.

### Validation of genes associated with relative chlorophyll, ΦPSII, and ΦNPQ using *Arabidopsis* T-DNA insertional mutants

Zm00001eb377130, a candidate gene linked to variation in chlorophyll content via GWAS, and one close paralog in the maize genome, Zm00001eb270460, are co-orthologous to a single *Arabidopsis* gene AT1G10500 (*ATCPISCA*). *Arabidopsis* plants homozygous for an insertion in *Atcpisca* exhibited a substantial reduction in chlorophyll content relative to wild type plants under both low light (200 µmol m^-2^ s^-1^) conditions and after 24 hours of high-light (550 µmol m^-2^ s^-1^) treatment (Fig. 4A). While this difference in phenotype was most significant when measured in vegetative stage plants (*p* = 0.0001), a similar pattern was observed in wild type and *Atcpisca* mutant plants at the flowering stage, with the difference being statistically significant under low light conditions (*p* = 0.006) but not statistically significant after high light treatment (Fig. S4). Zm00001eb386270, the sole candidate gene associated with the trait associated genetic marker for ΦPSII on chromosome 9 is orthologous to two duplicate genes in the Arabidopsis genome: AT2G26660 (*ATSPX2*) and AT5G20150 (*ATSPX1*). No significant differences were observed between *Arabidopsis* plants homozygous for an insertion in *Atspx2* and wild type plants under low light condition, but *Atspx2* mutant plants exhibited significantly higher ΦPSII than wild type plants after a 24 hours high-light treatment at both the vegetative (*p* < 0.0001) and flowering stages (*p* = 0.03) (Fig. 4B; Fig. S4). Zm00001eb378270, the most proximal of the five potential candidate genes associated with the trait associated genetic marker for ΦNPQ on chromosome 9 has a 1:1 orthologous relationship with AT5G42520 (*ATBPC6*). At the vegetative stage *Arabidopsis* plants homozygous for an insertion in *Atbpc6* exhibited substantially reduced ΦNPQ under both low light conditions and high light treatment (*p* = 0.0001), but no statistically significant differences between mutant and wild type plants were observed at the flowering stage (Fig. 4C; Fig. S4).

**Fig. 4.**
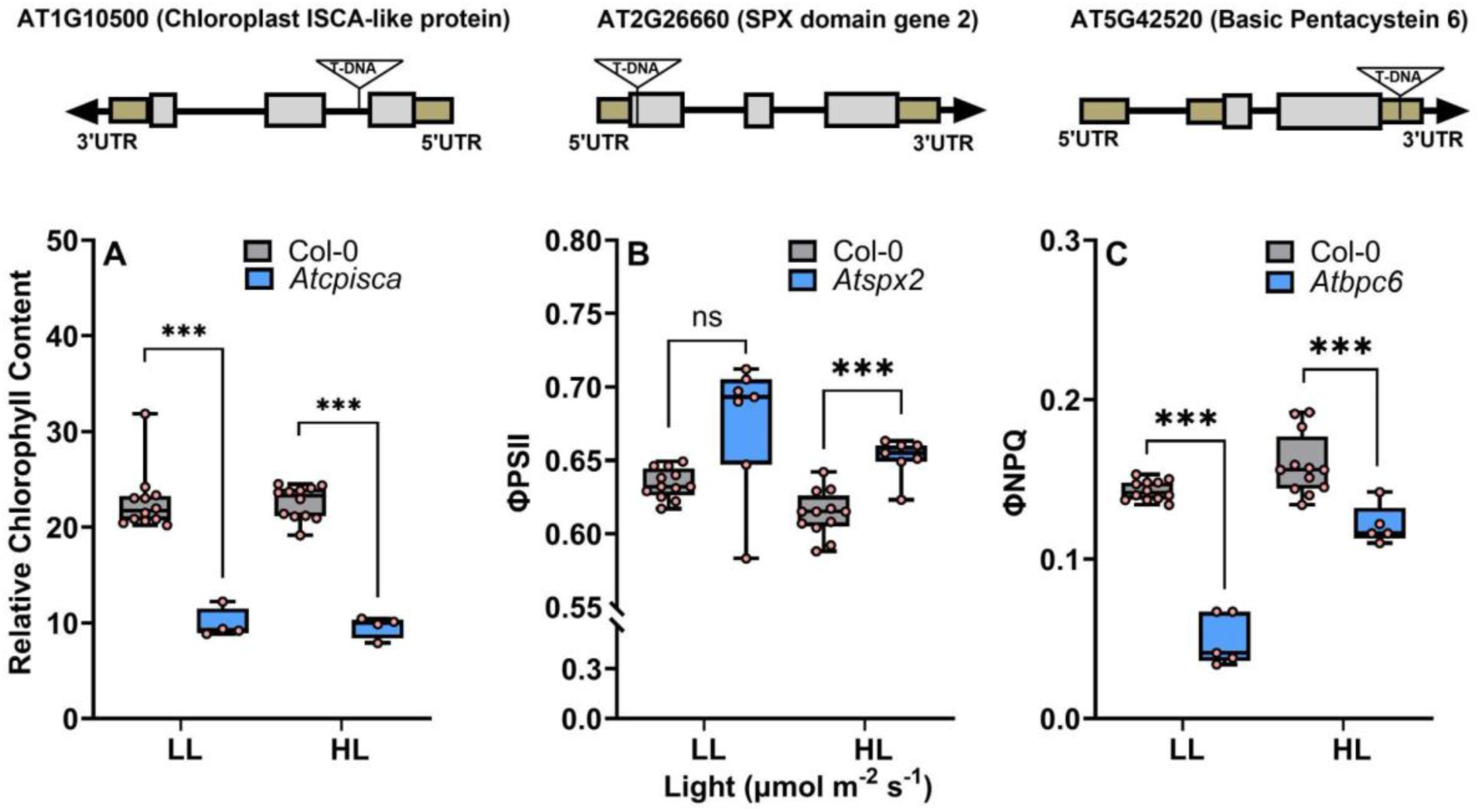
Phenotypes of insertion alleles of *Arabidopsis* genes homologous to maize candidate genes identified via GWAS. Plants were grown under low light conditions (LL; 200 µmol m-2 s-1) and moved to high light conditions (HL; 550 µmol m-2 s-1) after four weeks and phenotypes were collected under both low light and high light after a 24-hour acclimation to higher light intensity. **(A)** Difference in relative chlorophyll content between *Atcpisca* mutant (*Chloroplast-localized ISCA protein*; SALK_007055; n=4) and wild type Columbia plants (CS6000; Col-0; n=12) for AT1G10500. (**B)** Difference in ϕPSII between *Atspx2* mutant (*SPX domain gene 2*; SALK_007055; n=7) and wild type Columbia plants (Col-0; n=12) for AT2G26660. **(C)** Differences in ϕNPQ between *Atbpc6* mutant (*Basic Pentacysteine 6*; SALK_0047115; n=5) and wild type Columbia plants (Col-0; n=12) for AT5G42520. *p\** ≤ 0.05*, p*\*\* ≤ 0.01, *p*\*\*\* ≤ 0.001 (unpaired, two-tailed t-test).

## Discussion

Plant populations exhibit significant diversity in photosynthetic properties in addition to being highly responsive to environmental perturbations. Measurements of chlorophyll fluorescence are widely used to monitor and quantify the photosynthetic status of plants (Murchie and Lawson, 2013). Previous efforts to employ chlorophyll fluorescence to quantity genetic variation in photosynthetic parameters across large plant populations have either removed leaves or leaf disks from the field to collect measurements under controlled sets of environmental parameters (Ferguson *et al*., 2023; Sahay *et al*., 2023), or have made limited efforts to control for confounding factors via the incorporation of replicate or block effects (Dramadri *et al*., 2021; Liu *et al*., 2023). Across a wide enough range of conditions, many photosynthetic parameters exhibit non-linear responses to change in light intensity or other environmental factors, however we found that, for the range of environmental values observed during our field data collection, a relationship between environmental and photosynthetic parameters could be reasonably approximated with a linear model (Fig. S5).

Controlling for spatial and environmental covariates using linear models allowed us to substantially increase heritability (Fig. 1C) relative to other field studies conducted using similar approaches. Two previous studies employed the same instrument for field based phenotyping of photosynthetic traits, an analysis of 256 common bean (*Phaseolus vulgaris* L.) accessions (Dramadri *et al*., 2021) and an analysis of 225 rice (*Oryza Sativa* L.) accessions (Liu *et al*., 2023). Three of the fourteen traits we analyzed in this study were also represented in both of these previous studies: ΦPSII, ΦNPQ, and ΦNO. In all three cases, the heritability we achieved after correcting for both spatial and environmental confounders substantially exceeded those reported in previous studies which were unable to control for these confounders. We observed a heritability of 0.52 for ΦPSII in maize, substantially higher than the 0.15 reported in common bean or the 0.22 reported in rice. While, in principle, this increased heritability could also be attributed to greater genetically controlled diversity for ΦPSII in the maize population we examine related to the rice and common bean populations employed in these comparison studies, the heritability we observed for ΦPSII prior to controlling for spatial and environmental confounders (0.14; Fig. 1A) is similar to those reported in these other studies, suggesting the improvement is more likely attributable to the analyses method rather than differences in diversity between the populations studied. Similar patterns are present for ΦNPQ where the reported heritabilities were 0.14 in common bean, 0.35 in rice, and we observed 0.53 in maize subsequent to spatial and environmental correction and in ΦNO where the reported heritability was 0.08 in common bean, 0.18 in rice, and we had 0.32 in maize subsequent to spatial and environmental correction.

Out of the six strongly supported (RMIP > 0.2) trait associated SNPs identified as part of this study, two were located in regions of locally elevated linkage disequilibrium which resulted in large numbers of potential causal genes, and four were located in low linkage disequilibrium regions associated with only one or several potential causal genes (Fig. 2). We obtained homozygous insertion mutants for *Arabidopsis* orthologs of maize candidate genes in three of the four cases where a trait associated SNP tagged only one or several genes. In all three cases, including one candidate gene each from genome wide association studies conducted for relative chlorophyll, ΦPSII, and ΦNPQ, the *Arabidopsis* mutant phenotype was consistent with the predicted function from the maize GWAS study (Fig. 2, 4).

The gene Zm00001eb377130/AT1G10500, linked to chlorophyll content in maize via GWAS and in *Arabidopsis* via mutant analysis, is homologous to a well characterized bacteria gene *ISCA* which functions in the creation of iron-sulfur (Fe-S) complexes (Ayala-Castro *et al*., 2008). Fe-S complexes are involved in electron transfer reactions within chloroplasts, driving the redox processes necessary for synthesizing chlorophyll precursors (Abdel-Ghany *et al*., 2005)). Specifically, Fe-S cluster proteins facilitate enzymatic activities such as those in magnesium chelatase and other steps critical to the conversion of intermediates into functional chlorophyll molecules. In *Arabidopsis* the protein encoded by this gene has been shown to be chloroplast localized (Abdel-Ghany *et al*., 2005). Disruption of this gene in *Arabidopsis* via T-DNA insertion was associated with a substantial decline in relative chlorophyll content (Figure 4A; Fig. S4A) consistent with the predictions of the GWAS analysis conducted in maize.

An insertion allele of one of the two *Arabidopsis* orthologs of *ZmSPX1*, a gene linked to variation in ΦPSII via GWAS, exhibited a change in ΦPSII relative to wild type plants. The *Arabidopsis* insertion mutant (*ATSPX2*) exhibited an increase in ϕPSII (Figure 4B; Fig. S4B), consistent with mutant plants utilizing a larger proportion of light energy for productive photochemistry than wild-type plants. However, this apparent increase in photosynthetic productivity is consistent with a previous report that edited alleles of *ZmSPX1* in maize exhibit increases in productivity (Luo *et al*., 2024) suggesting *ZmSPX2* may act as a negative regulator of photosynthetic capacity. However caution should be taken in interpreting this result as many genes which apparently result in increases in yield or plant productivity fail to validate or exhibit significant phenotypic tradeoffs when tested across a wider range of environments (Khaipho-Burch *et al*., 2023).

The maize candidate gene *ZmBBR4* (Zm00001eb378270) belongs to a plant specific family of transcription factors with four members in maize and seven members in *Arabidopsis*. Analysis of both transcription factor binding site enrichment and coexpression networks suggest that *ZmBBR4* represses the expression of many photosynthesis related genes and contributes to both diurnal regulation of photosynthetic genes and the differentiation of expression between mesophyll and bundle sheath cells (Borba *et al*., 2023). Mutation of rice ortholog of *ZmBBR4*, *OsGBP1*, was associated with great biomass accumulation and larger seeds, with overexpression lines exhibiting reciprocal phenotypes (Gong *et al*., 2018). In *Arabidopsis*, single mutants in this transcription factor family had been reported to be phenotypically silent, while higher order mutants carrying loss of function alleles for multiple genes in this family did exhibit phenotypic effects (Monfared *et al*., 2011; Hecker *et al*., 2015). However, in our analysis, *Arabidopsis* plants carrying an insertion allele of *Atbpc6*, one of the *Arabidopsis* orthologs of *ZmBBR4*, exhibited decreases in ΦNPQ relative to wild type plants (Fig. 4C; Fig. S4C), a phenotype which was likely not screened for in prior efforts to characterize this gene family, demonstrating the potential of quantitative genetic analysis, even in distantly related species, to guide reverse genetics efforts in model species such as *Arabidopsis*.

The two GWAS hits which tagged high linkage disequilibrium windows in the maize genome each were associated with larger numbers of candidate genes (Fig. 2E, F; Table S4). While we did not validate any of these genes in these intervals via characterization of insertion alleles, each contains one or more genes with potential mechanistic links to variation in photosynthetic performance. The 22 annotated genes in the interval on maize Chromosome 8 associated with variation in ΦPSII include Zm00001eb348280, which encodes a member of the FtsZ2 protein family a group of proteins which play a role in chloroplast division. Two-fold overexpression of *AtFtsZ2-1* in *Arabidopsis* increased the number of chloroplasts whereas a three-fold increase reduces the number of chloroplasts (Stokes *et al*., 2000), while loss-of-function mutants of *AtFtsZ2-1* show severe defects in chloroplast division resulting in an increase in the size and a decrease in the number of chloroplasts in leaf mesophyll cells (Mcandrew *et al*., 2008; Schmitz *et al*., 2009). The same interval also contains Zm00001eb348290 which encodes a protein containing a 2Fe-2S ferredoxin-type iron-sulfur binding domain, particularly notable given the validated role of another gene, ISCA, associated with another GWAS hit from this same study which is involved in iron-sulfur binding. The interval on chromosome 3 associated with variation in ΦNPQ included 12 annotated genes. One of these, Zm00001eb140660, encoded a NAD(P)H-dependent oxidoreductase whose *Arabidopsis* ortholog, *AT3G2789*0 has been shown to play a role in regulating the production of reactive oxygen species (Biniek *et al*., 2017), a consequence of excess photosynthetic energy which is not safely dissipated via the NPQ pathway.

The successful functional validation of a number of the candidate genes identified via GWAS in maize using reverse genetics in *Arabidopsis* supports two assertions. Firstly, while maize and *Arabidopsis* belong to plant lineages separated by more than one hundred million years of evolution, the core components and regulators of photosynthesis related traits appear to be reasonably well conserved. This is consistent with several other recent cross species analyses of genes involved in photosynthetic trait variation (Sahay *et al*., 2023, 2024) and stands in contrast to the significant divergence on the roles of genes involved in determining phenotypes such as flowering time or plant morphology. Secondly, controlling for multiple confounding factors not only substantially increased heritability and substantially improved outcomes from genome wide association studies, resulting in both more and stronger GWAS hits. Collectively the results of this study provide reasons for optimism about the feasibility of studying the genetic determinants of variation in photosynthetic performance under field conditions and ultimately applying the insights gained to both engineering and breeding more photosynthetically productive crops.

## Supporting data

The following supplementary data are available at JXB online.

**Table S1** Range of values considered biological plausible and retained for quantitative genetic analyses for each trait analyzed in this study.

**Table S2** Primer sequences used for PCR to verify the *Arabidopsis* T-DNA insertional mutants.

**Table S3** Percentage of total trait variance explained by different experimental or environmental factors.

**Table S4** Absolute gene distance from genomic position of GWAS hit for mentioned traits

**Fig. S1.** Distribution of best unbiased linear estimates calculated after spatial correction for each of the fourteen traits employed in this study.

**Fig. S2.** Light intensity values recorded as part of photosynthetic trait phenotyping efforts.

**Fig. S3.** Results of a genome wide association study conducted using BLUEs calculated while correcting for device to device variation.

**Fig. S4.** Phenotypes of insertion alleles of *Arabidopsis* genes homologous to maize candidate genes identified via GWAS at flowering stage.

**Fig. S5.** Relationships between light intensity and photosynthetic trait values.

## Author contributions

JCS, RM, and MG designed and conducted the field experiment. MG and SS designed, organized and performed the collection of the photosynthetic data with input from JCS and RLR. WA conducted the quantitative genetic analyses with assistance and input from RM, JVTR, NS, FL and JCS. RKM contributed additional computational analysis. WA and SS conducted the *Arabidopsis* loss-of-function experiments and characterization with assistance from GB and guidance from RLR. WA and SS visualized the data with input from JCS. WA, SS and JCS interpreted the results with assistance and input from FL and RLR. WA, SS and JCS drafted the manuscript.

## Acknowledgments and funding

We thank Christine Smith for management of the field experiments, Annie Nelson, Rachel Gerdes, Bailey McLean, Aime Nishimwe, Michael Tross, and Mackenzie Zwiener for assistance in data collection. This research was supported by a UNL Wheat Innovation Fund grant to Rebecca Roston and James Schnable, a Higher Education Commission of Pakistan fellowship to Waqar Ali, a Foundation for Food and Agriculture award (602757) to James Schnable, and University of Nebraska-Lincoln startup funds to Seema Sahay.

## Data availability

Photosynthetic parameters measured as part of this study are provided as Supplementary Data File S1 and complete results from genome-wide association studies conducted here are provided as Supplementary Data File S2. https://figshare.com/s/8af0d240a1f39e6b6885. The genetic markers used in this study were subset from the file “*WiDiv.vcf.gz*” hosted on the Dryad repository: https://doi.org/10.5061/dryad.bnzs7h4f1 (Grzybowski et al., <u>2023</u>). Gene expression data employed in this study is publicly hosted on FigShare https://doi.org/10.6084/m9.figshare.24470758.v1 and has been previously described (Torres-Rodríguez *et al*., 2024). The code employed to conduct analyses and generate figures in this paper is deposited in an associated GitHub repository: https://github.com/waqarali1994/MultispeQ.git

